# Microbicidal Mechanisms for Light-Activated Molecular Nanomachines in *Mycobacterium smegmatis*: A Model for Pathogenic Bacteria

**DOI:** 10.1101/2024.10.04.616754

**Authors:** Thushara Galbadage, Dongdong Liu, James M. Tour, Jeffrey D. Cirillo, Richard S. Gunasekera

## Abstract

There is a global health crisis of antimicrobial resistance, with over a million deaths annually attributed to antimicrobial-resistant pathogens, and mycobacterial infections are a major cause of antimicrobial-resistant infections, leading to more deaths than any other single infectious agent. Notably, the rise of multidrug-resistant (MDR), extensively drug-resistant (XDR), and totally drug-resistant (TDR) strains of *Mycobacterium tuberculosis* led to higher mortality rates and challenge all existing antibiotic regimens. Light-activated molecular nanomachines (MNMs) represent a promising class of broad-spectrum antimicrobial agents that could help counter this rise in antimicrobial resistance. Addressing a key knowledge gap, this study explores the mechanisms of action for MNMs in *Mycobacterium smegmatis*, a surrogate model for pathogenic mycobacteria. We show that fast rotor MNMs kill up to 97% of *M. smegmatis* and co-localize with the bacteria as part of their mechanism of action. The ability to translate these observations to pathogenic mycobacteria was demonstrated by the ability of MNMs to kill 93.5% of *M. tuberculosis* under similar conditions. These findings suggest that MNMs may provide innovative sustainable antimicrobial agents for the treatment of drug-resistant mycobacterial infections.

**Graphical Abstract:** Bacteria exposed to MNMs have two distinct outcomes when activated by 365 nm light. Slow motors (MNM **2** and **4)** have no rotational action, remains outside the bacteria and have little to no effect on bacterial viability. Whereas fast motors (MNM **1** and **3)** co-localize and embed into the bacterial cell wall causing disruptions that lead to a significant reduction in bacterial viability.

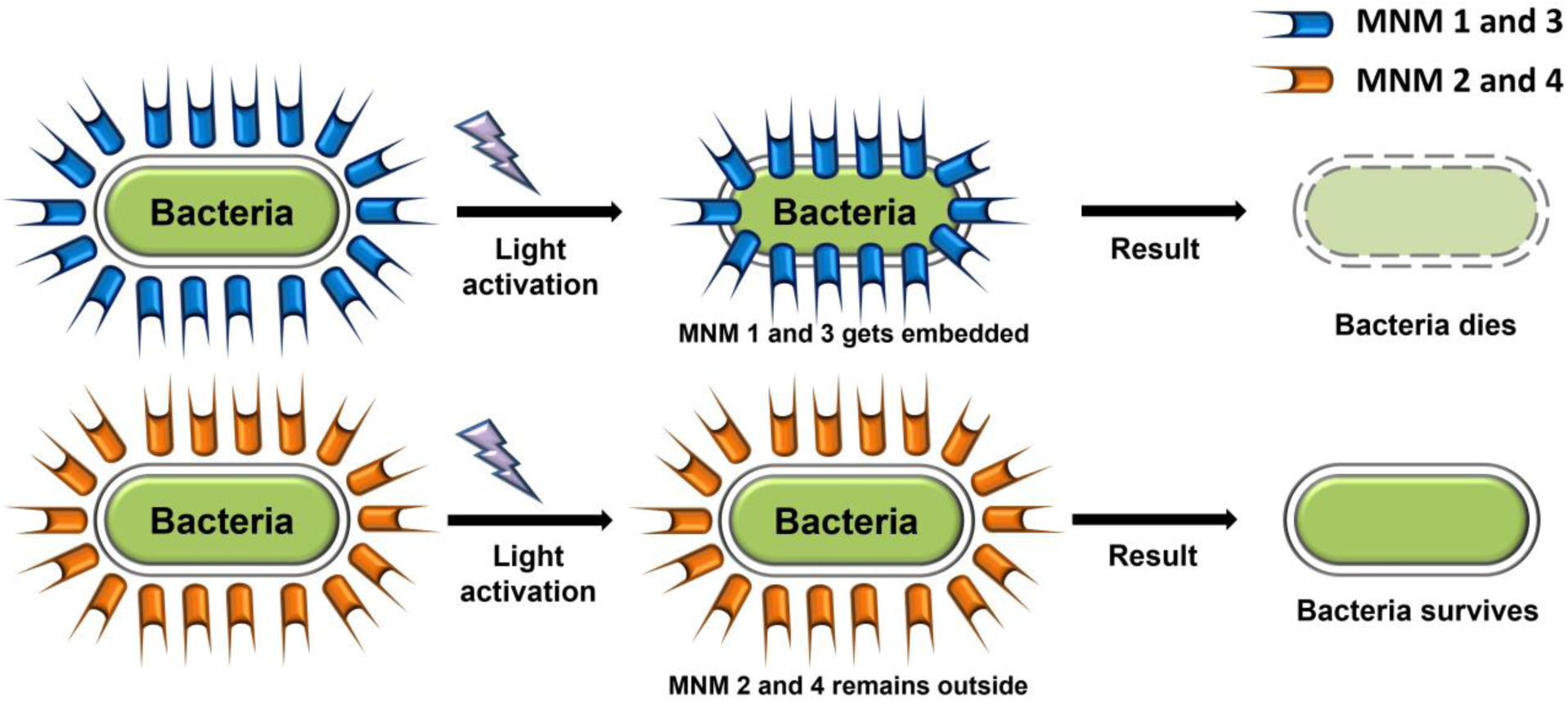

## Introduction

Antimicrobial resistance (AR) is an increasing global health crisis with profound implications for medical and public health worldwide (1, 2). The persistence and rise of AR pose a significant public health threat, as evidenced by over a million deaths annually attributed to antimicrobial-resistant (AMR) pathogens (1). If current rates of AR continue, it is projected that over 10 million annual deaths will be caused by AMR pathogens by 2050 (3).

Mycobacterial infections are one such challenging global threat, with several species of mycobacterial pathogens emerging as significant concerns, particularly due to the growing incidence of antibiotic resistance and multidrug-resistant strains (4–6). Especially, *Mycobacterium tuberculosis*, the causative agent of tuberculosis, and *Mycobacterium leprae*, responsible for leprosy, are important pathogens causing chronic and debilitating infections in humans. The ability of these bacteria to develop resistance to multiple drugs poses a serious obstacle to currently effective treatment, necessitating prolonged and often complex therapeutic regimens (7). Multidrug-resistant (MDR), extensively drug-resistant (XDR), and totally drug-resistant (TDR) strains of *Mycobacterium tuberculosis* are some of the most important public health problems worldwide, and perhaps even more important than ESKAPE pathogens (7–9). MDR, XDR, and TDR lead to higher mortality rates and challenge all existing antibiotic regimens (7, 10). The rapid development of MDR, XDR, and TDR strains further exacerbates the difficulty in managing mycobacterial infections, making it imperative to address antibiotic resistance comprehensively.

There are three main mechanisms that pathogens use to develop antimicrobial resistance. First, the accumulation of various resistance genes or pathogenicity islands through mobilization and horizontal gene transfer from various environmental bacteria. Second, random mutants that occur in antibiotic target genes that make them have fewer active sites for antibiotic action. Third, upregulation of inherent resistance mechanisms, including antibiotic inactivating enzymes and efflux system to pump out antibiotics that enter the pathogen (11, 12). These forms of antimicrobial resistance mechanisms can render all conventional and targeted antimicrobial agents ineffective over time. Therefore, innovative antimicrobial agents that can circumvent antimicrobial resistance mechanisms must be developed.

To this end, our collaborative research group has characterized molecular nanomachines (MNMs) as a unique class of nanomechanical broad-spectrum antimicrobial agents that have the potential to remain effective long-term (13, 14). MNMs are synthetic organic nanomolecules that can rotate unidirectionally up to several million rotations per second upon light-activation (Figure 1) (15–18). Our approach using MNMs bypasses the need for specific pathogen interactions and likely renders the pathogen incapable of acquiring resistance to these MNM agents. More recently, MNMs were shown to effectively eliminate antibiotic-resistant Gram-negative and Gram-positive bacteria, including methicillin-resistant *Staphylococcus aureus*, while preventing resistance development in an *in vivo* model of burn wound infection (14).

**Figure 1.**
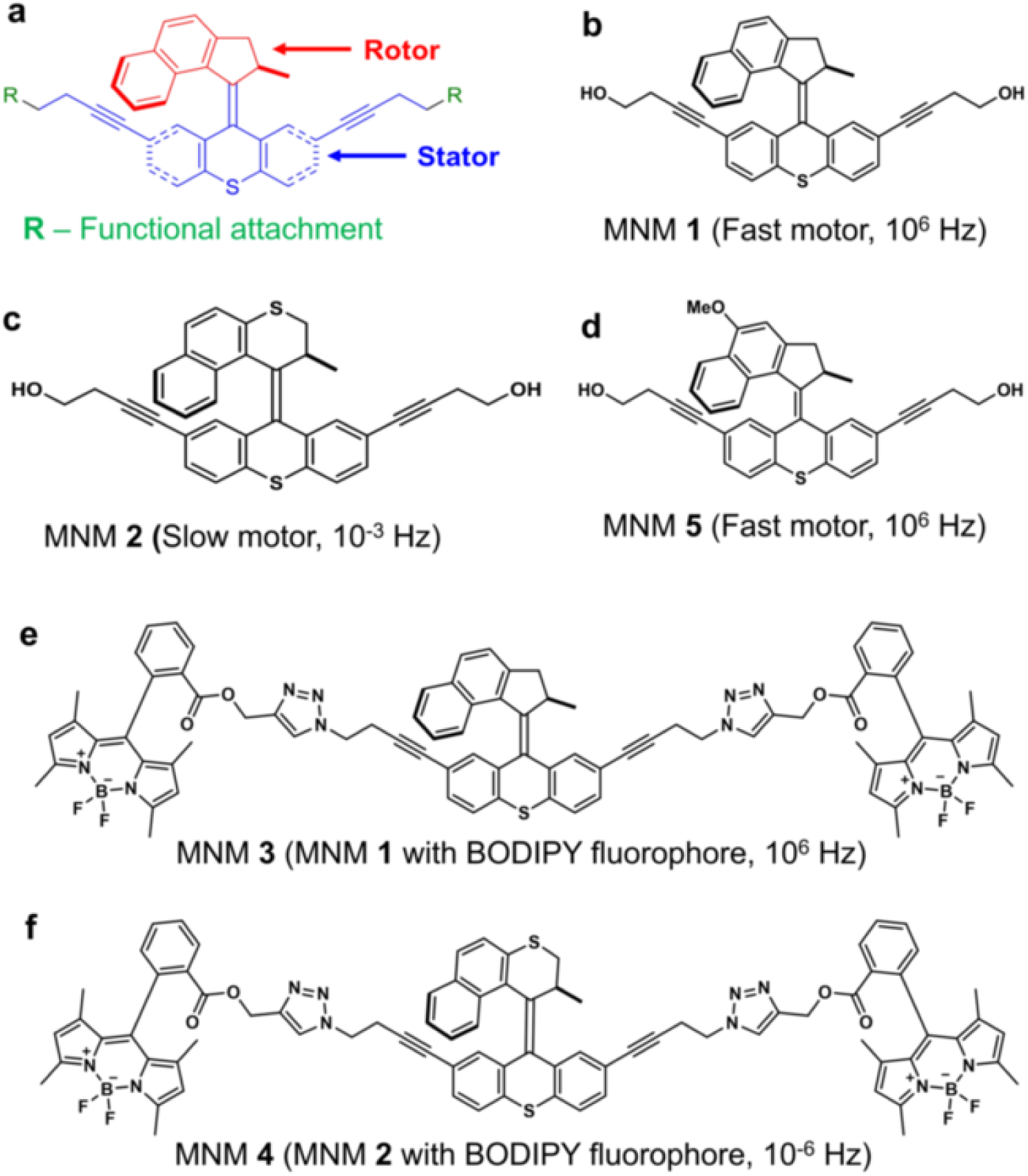
Molecular nanomachines (MNMs) were used in this study. (a) A graphical representation of the MNM structure showing the rotor and stator components. (b) MNM **1** is a fast motor with a rotor that can rotate at 2-3 MHz when activated with 365 nm light. (c) MNM **2** is a slow motor with a rotor that can rotate a 1.8 revolutions per hour when activated with 365 nm light. (d) MNM **5** is a fast motor with a rotor that can rotate at 2-3 MHz when activated with 395 nm light. (e) MNM **3** fast Motor with BODIPY fluorophore attached to two arms in its stationary component and activated by 365 nm light. (f) MNM **4** slow motor with BODIPY fluorophore attached to two arms in its stationary component and activated by 365 nm light.

In recent studies, the mechanism of action of MNMs has been explored, focusing on the physical membrane disruption and damage done to bacterial or eukaryotic cells. Electron microscopy (EM), membrane permeability assays, and RNA-seq methods were used to characterize the effects of MNMs on bacterial cells (13, 14). While these methods illustrated various aspects of MNM activity, the mechanism of action of these MNMs is not yet fully understood, posing an important knowledge gap that needs to be addressed. Here we use *Mycobacterium smegmatis* as a model for mycobacterial species to show that light-activated MNMs can act as antibacterial agents against mycobacteria, reduce bacterial viability, and further characterize the mechanism of action of MNMs. *Mycobacterium tuberculosis*, *Staphylococcus aureus*, and *Escherichia coli* were used as relevant pathogenic mycobacterial species, a Gram-positive control and a Gram-negative control, respectively.

*M. smegmatis* is an acid-fast staining bacterium with a lipid-rich cell wall containing mycolic acid (19). While *M. smegmatis* are non-pathogenic free-living mycobacteria, there are several related pathogenic mycobacteria, such as *M. tuberculosis, M. avium,* and *M. leprea* that are of significant clinical importance (20). Tuberculosis caused by *M. tuberculosis* is the leading cause of death worldwide due to a single infectious agent and the ninth leading cause of death overall (21). *M tuberculosis* has multidrug-resistant (MDR) and extremely drug-resistant (XDR) strains that have significant implications in immune-compromised patient populations (22). *M. smegmatis* is a relevant and commonly used mycobacterium model to study cellular processes that are related to the *M. tuberculosis* pathogen (20).

In this study, we exposed *M. smegmatis* to light-activated MNMs, observed the reduction in bacterial viability, and characterized their nanomechanical action using fluorescently labeled MNMs, and confocal microscopy. Here we show that fast motor MNMs can reduce mycobacterial viability, with their nanomechanical action by co-localizing with the bacteria. Our study seeks to further explore the use of MNMs to counter antibiotic resistance, safeguard public health, and develop innovative interventions for a sustainable and resilient future in the face of the formidable global health threat, of antibiotic resistance.

## Methods

### Bacterial strains

A wild-type *Mycobacterium smegmatis* (mc^2^155), mutant *Mycobacterium tuberculosis* (mc^2^7000), bioluminescent strain of *Staphylococcus aureus* (Xen36) (ATCC 49525:: luxABCDE operon), and wild-type *E. coli* (k-12 strain, HB101) were used to characterize the bactericidal properties of MNMs against these bacterial species. A tdTomato fluorescent *M. smegmatis* strain (ψms23) was derived by transforming wild-type a wild-type *M. smegmatis* (mc^2^155) with a multi-copy plasmid, pJDC60 (pFJS8ΔGFP::tdTomato, under a PL5 promoter, with kanamycin selection). The fluorescent strain of *M. smegmatis* was used for confocal microscope imaging to explore the co-localization of MNMs and bacteria.

### Molecular nanomachines

MNMs are molecular motors that have a stator and a rotor component, with different functional groups attached to the stator (Figure 1 a). MNM **1** is a fast motor that rotates at about 2-3 x 10^6^ revolutions per second (10^6^ Hz) (Figure 1 b). MNM **2** is a slow motor that rotates about 1.8 revolutions per hour (10^-3^ Hz) (Figure 1 c). MNM **2** served as a non-rotating analog to MNM **1**. MNM **3** and MNM **4** are MNM **1** and MNM **2** respectively, with two BODIPY fluorophores attached to the stator (Figure 1 e and f). MNMs **3** and **4** were used for used for confocal microscope imaging. MNMs **1** to **4** are activated with 365 nm light. MNM 5 is a fast motor that rotates at about 2-3 x 10^6^ revolutions per second (10^6^ Hz) and is activated with 395 nm light (Figure 1 d, Supplemental Figure 1).

### Bacterial viability assays with 365 nm light-activated MNMs

Overnight cultures of bacteria grown in either Luria-Bertani (LB) or Middlebrook with ADC (MADC) media were used to start secondary cultures. Cultures were placed in a shaker at 37 °C for about 4 hours to obtain fresh bacterial cultures in the log phase (Figure 2 a). Serial dilutions of cultures were made using 100 µL of culture with 900 µL of 1x PBS. Two 1 ml dilutions of 10^-4^ or 10^-5^ cultures were used for the viability assays. The concentration of MNMs in 1% DMSO was either 1 or 10 µM. 1 µM concentration of MNMs was used for *M. smegmatis*, *M. tuberculosis*, and *S. aureus*. 10 µM concentration of MNMs was used for *E. coli*. Using 10 mM working stock of MNM, 1 µL was added to 1 ml of bacterial culture at 10^-5^ to give a 10 µM concentration. 1 µL of DMSO without MNM was added to the second 1 ml bacterial culture at 10^-5^ as the control (No MNM control). They were incubated for 30 minutes on a test tube rocker at room temperature. At 30 minutes after incubation, 120 µl of the culture of each was transferred onto two petri dish lids each and one of each (DMSO control, slow motor, and fast motor) was exposed to 365 nm light (UV-c) for 5 minutes (Figure 2 b and c). The distance between the light source and the bacterial culture was 1.3 cm (0.5 inches) (Figure 2 d). The other set was not exposed to light and was used as the non-activated control. After light exposure, the bacterial cultures were serially diluted and plated on 60 x 15 mm Petri dishes with LB or MADC agar and were incubated at 37 °C overnight for growth. Colonies were counted and the bactericidal effect of MNM on the different bacterial species was determined.

**Figure 2.**
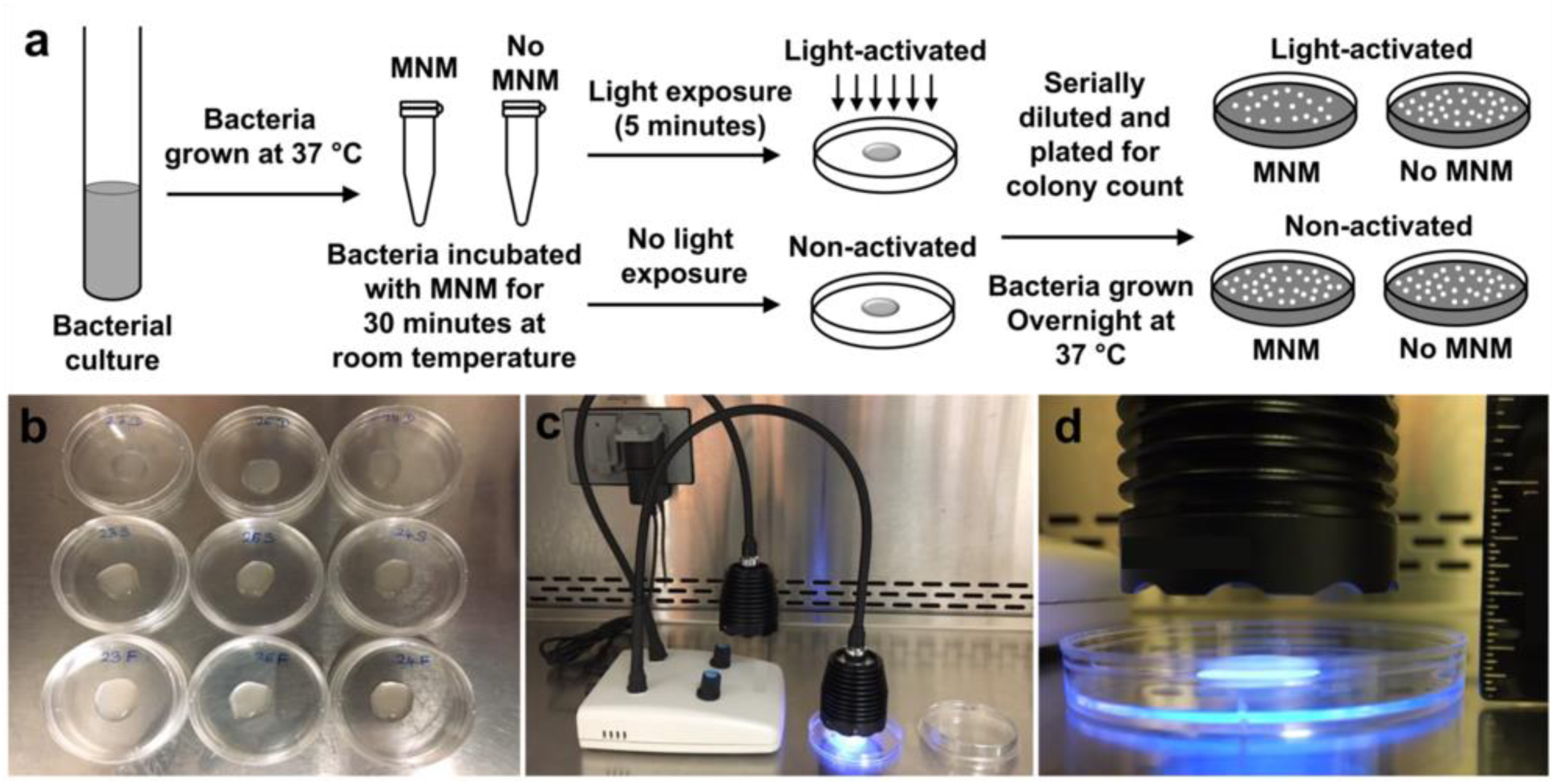
Bacterial viability assay with light-activated molecular nanomachines (MNMs) Setup. (a) Experimental setup for bacterial viability reduction assays. A log growth phase bacterial culture was incubated with no MNM (with dimethyl sulfoxide, DMSO), MNM **2**, or MNM **1** for 30 min, activated with 365 nm light for 5 min, and plated for CFU/mL counts. (b) 120 µl of bacterial culture was placed on the lid of a petri dish for light exposure. (c) 365 nm light source used to activate MNMs **1** to **4**. Bacterial cultures were exposed to 365 nm light for 5 minutes. (d) The light source was placed directly above the bacterial culture at a constant distance from the culture (1.3 cm or 0.5 inches), to ensure uniform and constant energy delivery.

### Bacterial viability assays with 395 nm light-activated MNMs

Using an overnight culture of wild-type *M. smegmatis* (mc2 155 strain), a secondary culture was started in a 4 ml volume. OD600 of the overnight culture was measured. The starting OD of the secondary culture was 0.1. This culture was placed at 37 C for about 3 hours to give an active culture of *M. smegmatis*. Ending OD600 of the secondary culture was measured. A serial dilution of this culture was made using 100 µl of culture with 900 µl in MADC media. 1 ml dilutions of 10^-4 cultures were used for the experiment. 1 µM concentration of MNMs in DMSO was used. Using a 10 mM working stock of MNMs, a 1 mM solution of MNMs was prepared with 2 µl of 10 mM MNMs with 18 µl of 1x PBS. 1 µl of this 1 mM MNMs was added to 1 ml of *M. smegmatis* culture at 10^-3^ to give a 1 µM concentration. 1 µl of DMSO without MNMs was added to the second 1 ml *M. smegmatis* culture at 10^-3^ as the control. The DMSO concentration for the experiment was 0.1%. They were incubated for 30 min on a test tube rocker at room temperature. At 30 min after incubation, 100 µl of the culture of each was transferred into a 96-well plate with 4 replicates each. These were exposed to 395 nm light for 0, 5, 15, and 30 min. The light source was directly plated on top of the 96-well plate. After 395 nm light exposure, serial dilutions of cultures were made and plated for CFU counts. These plates were incubated at 37 ^C^ overnight for colony growth.

### Confocal microscopy to observe MNM and *M. smegmatis* interactions

A confocal microscope (Nikon A1R+) with spectral capability, lasers FITC (fluorescein isothiocyanate) with an excitation wavelength of 488 nm and TRITC (tetramethylrhodamine isothiocyanate) excitation wavelength of 561 nm was used to detect excitation of BODIPY (505– 512 nm) and tdTomato (575–585 nm), respectively. Using a 60× oil immersion objective, ψms23 (*M. smegmatis*::tdTomato) were imaged 30 minutes post-exposure to MNM **3** and MNM **4** tagged with BODIPY. After exposure to MNM, ψms23 was washed with 1x PBS, fixed with 4% paraformaldehyde, and washed once after fixation. 10 µl of No MNM, MNM **3,** and MNM **4** exposed ψms23 were then mounted on slides with 8 µl of mountant solution each. They were each imaged at 8 different visual fields at a magnification of 600x and a resolution of 1024. Representative images are presented to show differences observed after exposure to non-activated and activated MNM.

### MNM and *M. smegmatis* fluorescent intensity quantification

NIH ImageJ software was used to quantify the fluorescent intensities of images obtained by confocal imaging. Confocal images were taken with each laser (TRIC and FITC) individually and a merged image. Quantification was done on each of the wavelengths 488 nm and 561 nm and of the merged image. For each group, 8 images taken were quantified using the same area, and averages were calculated. For each group, the ratios of the MNM to *M. smegmatis* were obtained and their averages were calculated. The results presented are an average of 8 separate images taken for each group.

### *In vivo* imaging system (IVIS) and quantification

IVIS Lumina II was used to quantify the total fluorescence of plates used for plating ψms23 post the day after carrying out viability assays. Each of the plates was imaged using an emission wavelength of 580 nm and an excitation wavelength of 535 nm. The IVIS settings were epi-illumination, Bin 4 (medium), FOV: 12.5, f2 and exposure time of 0.5 s. Three replicates for each group were imaged. Living Image software was used to analyze and quantify the fluorescence intensity. A region of interest (ROI) was selected to include the whole plate and background fluorescence was reduced from each image to normalize readings from each plate. The units of measurement were radiance (photons) with units of p/sec/cm^2^/sr. All plates used for quantification are included in the image, with lower and upper limits of 4.5 x 10^7^ and 1.0 x 10^8^ respectively.

### Statistical analyses

All experiments were carried out with at least an n of 3. The numbers of replicates used in each experiment are stated in each experiment if more than 3 were used. GraphPad Prism was used to perform two-tailed unpaired t-test statistical analyses to compare the means of different bacterial exposure groups. Means and standard errors are presented in each of the graphs plotted in Microsoft Excel.

## Results

### Light-activated MNM 1 reduces the viability of *M. smegmatis*, *M. tuberculosis*, *S. aureus* and *E. coli*

We exposed *M. smegmatis* (mc^2^ 155) to 1 µM of MNM **1** with 5 minutes of 365 nm light-activation and observed a 40% relative reduction in bacterial viability compared to a non-activated MNM **1** control (Figure 3 a and b). This reduction in bacterial viability is attributed to the action of activated MNM **1**, as 365 nm light alone, No MNM, MNM **2,** or non-activated MNM **1** showed little deleterious effect on *M. smegmatis* (≤ 10% viability reduction). Similarly, we exposed *M. tuberculosis* (mc^2^7000) to 1 µM of MNM **1** with 5 minutes of 365 nm light-activation and observed a 93% relative reduction in bacterial viability compared to a non-activated MNM **1** control (Figure 3 c and d). However, *M. tuberculosis* was much more sensitive to the deleterious effects of exposure to 5 minutes of 365 nm light, as both the No MNM and MNM **2** controls showed 70% and 75% viability reduction upon light exposure. This observation confirms previous findings that *M. tuberculosis* is sensitive to light and that UV light can be used to control the environmental spread of pathogenic mycobacteria (23–25). Within this context light-activated MNM **1** still reduced *M. tuberculosis* significantly compared to No MNM (p = 0.0102) and MNM **2** (p = 0.0078). This shows that light-activated MNM **1** can significantly reduce the viability of both *M. smegmatis* and *M. tuberculosis*.

**Figure 3.**
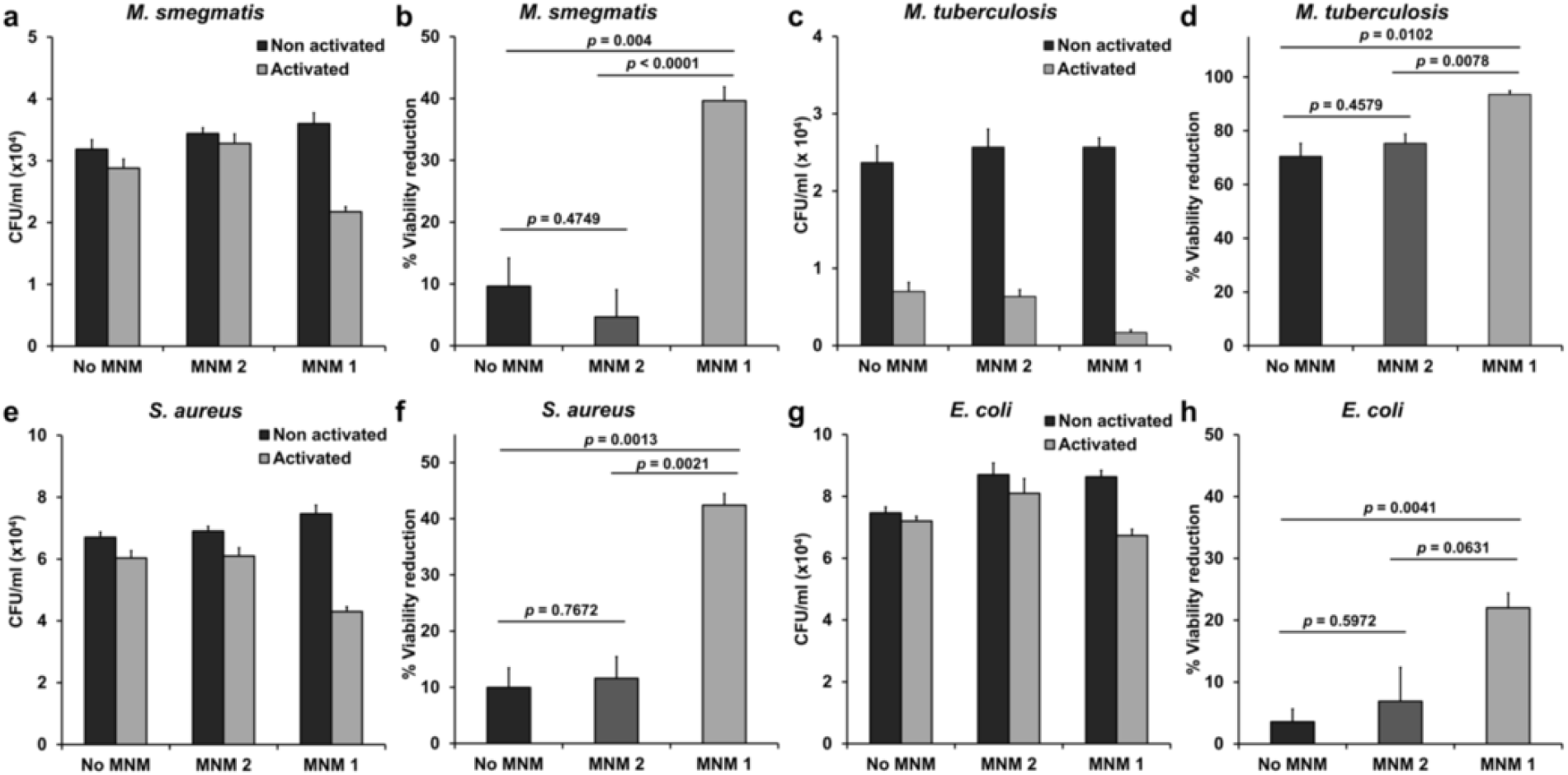
Viability reduction of bacteria with 365 nm light-activated molecular nanomachines (MNMs). Four different bacterial species were exposed to MNMs with 5 minutes of 365 nm light at 14 mW/cm^2^ (activated) and compared to the same samples not exposed to light (non-activated). No MNM is a 0.1% DMSO control. MNM **2** is the slow motor. MNM **1** is the fast motor. (a) *M. smegmatis* (mc^2^ 155) and 1 µM of MNMs with or without 5 min of 365 nm light activation. (b) Light-activated No MNM, MNM **2** and MNM **1** groups show 10%, 5%, and 40% reduction in viability of *M. smegmatis* compared to the non-activated controls, respectively. (c) *M. tuberculosis* (mc^2^7000) and 1 µM of MNMs with or without 5 min of 365 nm light activation. (d) Light-activated No MNM, MNM **2** and MNM **1** groups show 70%, 75%, and 93% reduction in viability of *M. tuberculosis* compared to the non-activated controls, respectively. (e) *S. aureus* (Xen 36) and 1 µM of MNMs with or without 5 min of 365 nm light activation. (f) Light-activated No MNM, MNM **2** and MNM **1** groups show a 10%, 12%, and 42% reduction in the viability of *S. aureus* compared to the non-activated controls, respectively. (g) *E. coli* (K-12) and 1 µM of MNMs with or without 5 min of 365 nm light activation. (h) Light-activated No MNM, MNM **2** and MNM **1** groups show 4%, 7%, and 22% reduction in viability of *E. coli* compared to the non-activated controls, respectively. Error bars represent the standard errors of the mean.

Mycobacteria are acid-fast bacteria and have a thick waxy layer on their cell wall that helps protect them from antibiotics and the host immune system (26). We were interested in assaying the bactericidal properties of light-activated MNM **1** against Gram-positive and Gram-negative bacteria. Therefore, we exposed *S. aureus* (Gram-positive) and *E. coli* (Gram-negative) to MNM **1**. When *S. aureus* (Xen 36) was exposed to 1 µM of MNM **1** with 5 minutes of 365 nm light-activation and observed a 42% relative reduction in bacterial viability compared to a non-activated MNM **1** control (Figure 3 e and f). No MNM, MNM **2** or non-activated MNM **1** showed little deleterious effect on *S. aureus* (≤ 12% viability reduction). This viability reduction was similar to what we observed with *M. smegmatis*. When *E. coli* was exposed to 10 µM of MNM **1** with 5 minutes of 365 nm light activation we observed a 22% relative reduction in viability compared to a control without MNM **1** (Figure 3 g and h). No MNM, MNM **2** or non-activated MNM **1** showed little deleterious effect on *E. coli* (≤ 7% viability reduction). Compared to *M. smegmatis* or *S. aureus*, *E. coli* showed relatively less viability reduction with light-activated MNM **1** (22% vs 40%). This could likely due to the cell wall structure of Gram-negative bacteria compared to Gram-positive or acid-fast bacteria and is similar to what we have observed with K. pneumoniae, another Gram-negative bacteria (13). These results show that 5-minute light activation of MNM **1** caused significant viability reductions in *M. smegmatis*, *M. tuberculosis*, *S. aureus* and *E. coli*, and have the potential to be used as a broad-spectrum antibacterial agent.

### Light-activated MNM 5 reduces viability in *M. smegmatis* by over 95%

Light-activated MNM **1** reduced *M. smegmatis* viability by 40% within 5 minutes. However, 365 nm light showed deleterious effects of *M. smegmatis* with prolonged exposure (Supplemental Figure 2 a). Therefore, we were not able to reasonability assess the bactericidal potential of MNM 1 with prolonged exposure without reducing bacterial viability with the 365 nm light. To address this, we used a different fast motor, MNM 5, that is activated with 395 nm light (Figure 1 d). Using a 395 nm light source that emitted 56.5 mW/cm^2^ of flux, we assess the deleterious effect of this source on *M. smegmatis* (Figure 4 a). Compared to the 365 nm light source, the 395 nm light had a smaller deleterious effect on *M. smegmatis* over one hour, reducing viability by 14% at 30 minutes and 35% at 60 minutes of light exposure (Figure 4 b and Supplemental Figure 2). Therefore, we exposed *M. smegmatis* to 1 µM of MNM **5** and activated it for a period of 30 minutes, assessing the viability reduction of *M. smegmatis* at 5, 10, 15 and 30 minutes (Figure 4 c and d). *M. smegmatis* was also exposed to 395 nm light without MNM 5 (No MNM) as a control. *M. smegmatis* percent viability reduction due to light-activated MNM 5 at 5, 10, 15 and 30 minutes were 8%, 21%, 35%, and 97%, respectively. At 15 and 30 minutes of light-activated MNM **5** showed a significant decrease in viability compared to the No MNM control with *p*-value = 0.0354 and *p*-value < 0.0001, respectively. These results show that fast motor MNM **5** was able to reduce *M. smegmatis* viability by 97% and indicate its potential as a clinically relevant antibiotic agent.

**Figure 4.**
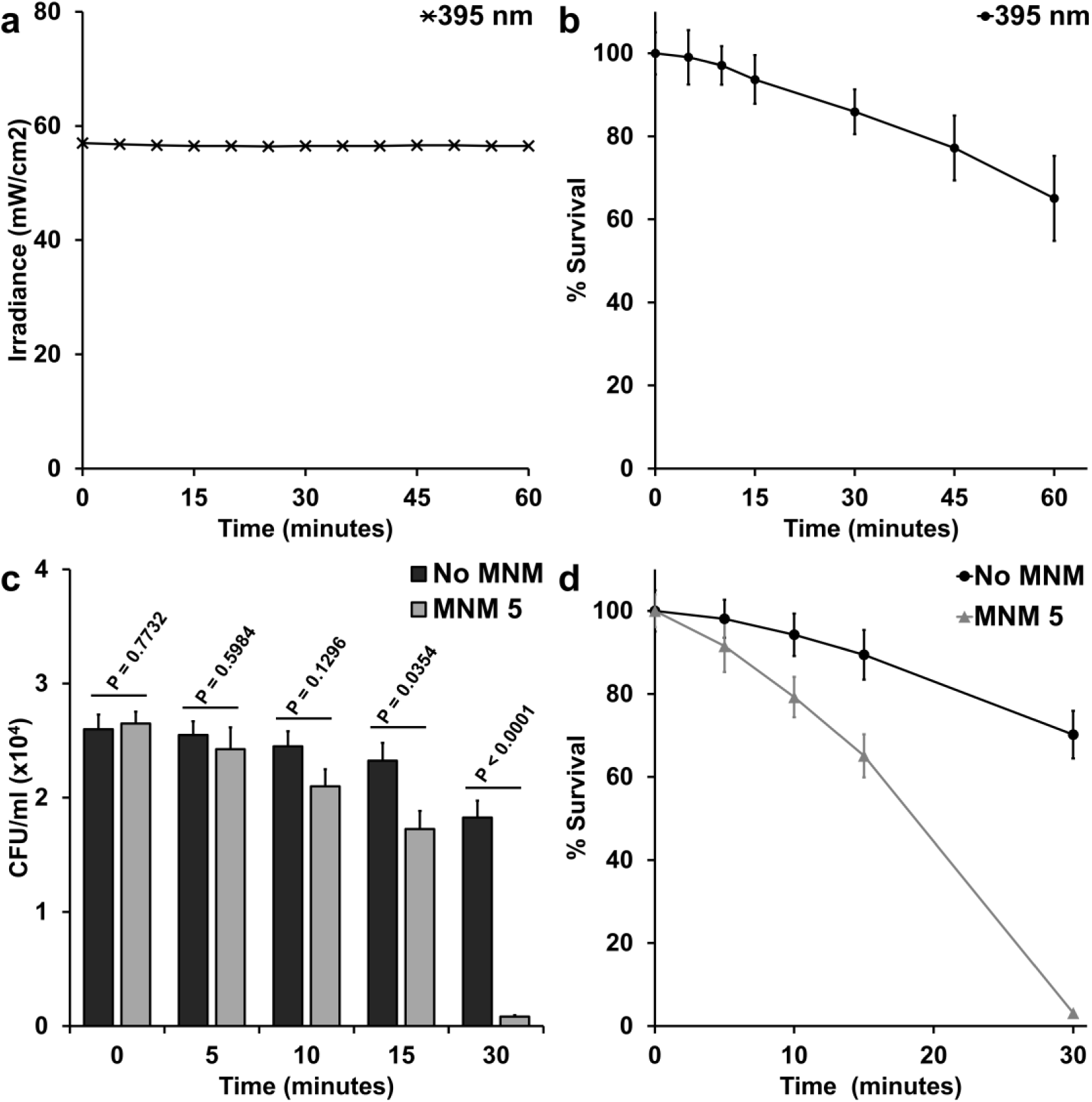
Viability reduction of *M. smegmatis* with 395 nm light-activated molecular nanomachines (MNMs). *M. smegmatis* (mc^2^ 155) was exposed to 1 µM of MNM **5** activated at different lengths of time to a 395 nm light source at 56.5 mW/cm^2^ to observe the reduction in bacterial viability caused by activated MNM **5**. (a) The irradiance of the 395 nm light source measured over one hour held steady within a range of 56.4 to 56.8 mW/cm^2^. (b) Bactericidal effects of the 395 nm light source at 56.5 mW/cm^2^ were observed over one hour. *M. smegmatis* percent viability compared to the starting culture at different lengths of light exposure: 5 minutes (99%), 10 minutes (97%), 15 minutes (94%), 30 minutes (86%), 45 minutes (77%), 60 minutes (65%). (c) Viability reduction of *M. smegmatis* exposed to light-activated MNM **5** compared to a No MNM control exposed to the same duration of 395 nm light. At 15 and 30 minutes of light-activated MNM **5** showed a significant decrease in viability compared to the No MNM control. (d) Bactericidal effects of light-activated MNM **5**. *M. smegmatis* percent viability compared to the starting culture at different lengths of exposure to light-activated MNM **5**: 5 minutes (92%), 10 minutes (79%), 15 minutes (65%), 30 minutes (3%). Error bars represent the standard errors of the mean.

### Light-activated fast motor MNM co-localizes with *M. smegmatis*

After characterizing the nanomechanical antimicrobial properties of light-activated MNM **1** and **5**, we were interested in studying the mechanism of action of these MNMs in *M. smegmatis* bacteria. For this we used MNM **1** and MNM **2** with fluorophore BODIPY attached to them forming MNM **3** and MNM **4**, respectively (Figure 1 e and f). *M. smegmatis* strain expressing tdTomato fluorophore (ψms23) was exposed to light-activated MNM **3** and **4** and imaged with confocal microscopy (Figure 5). We observed a 3x increase in fluorescent intensity in MNM- BODIPY levels in ψms23 exposed to light-activated MNM **3** compared to non-activated MNM **3** (p-value = 0.0065) (Figure 5 a-f and s). In contrast, there was no change in fluorescent intensity in MNM-BODIPY levels in ψms23 exposed to non-activated or light-activated MNM **4** (p-value = 0.4106) (Figure 5 g-l and s). Control ψms23 imaged without any MNM-BODIPY showed background fluorescent levels were minimal (Figure 5 m-r).

**Figure 5.**
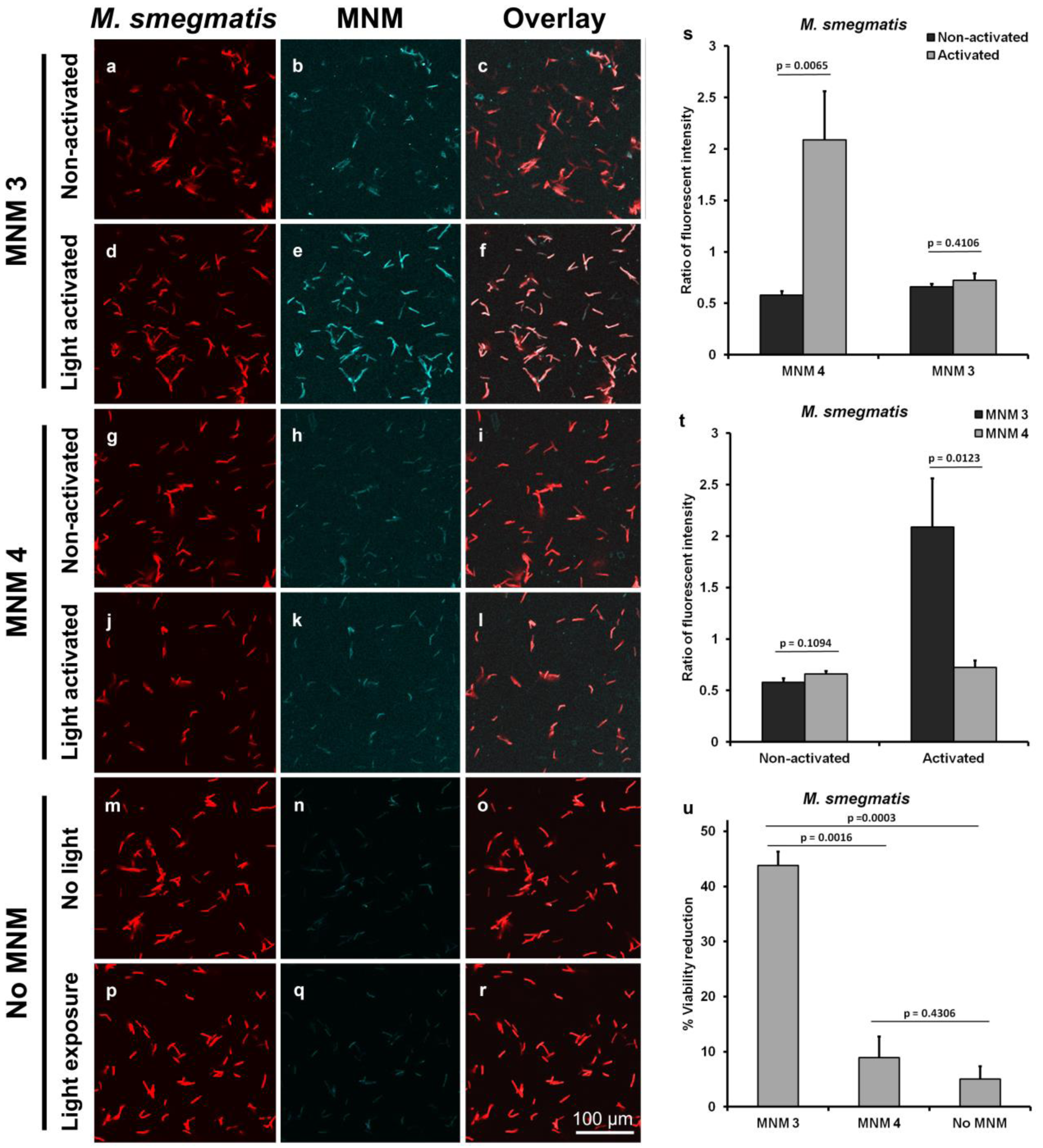
High-resolution confocal images show light-activated MNM 3 co-localize with *M. smegmatis.* *M. smegmatis* expressing tdTomato (ψms23) were exposed to 10 µM of fluorescent MNM-BODIPY and were imaged at 30 minutes after exposure to non-activated or light-activated MNM. Bacteria with MNM **3**, MNM **4,** or without MNM activated with 365 nm light for 5 minutes and compared with the non-activated group. A 60x oil-immersion objective was used with excitation wavelengths of 488 for BODIPY and 561 nm for tdTomato. (a-c) Non-activated MNM **3**. (d-f) Light-activated MNM **3**. (g-i) Non-activated MNM **4**. (j-l) Light-activated MNM 4. (m-o) No MNM control without light exposure. (p-r) No MNM control with light exposure. Column *M. smegmatis* shows the bacterial imaged at 561 nm. Column MNM shows the BODIPY-MNM images at 488 nm. The overlay column shows the overlay of the *M. smegmatis* image and the MNM images for each group. (s-t) Confocal imaging quantification using ImageJ. (s) Comparison of MNM **3** and MNM **4** co-localized *M. smegmatis*. (t) Comparison of co-localization of MNM with and without light activation (u) tdTomato expressing *M. smegmatis* (ψms23) exposed to 10 µM of MNM. The relative reduction in viability of *M. smegmatis* exposed to light-activated MNM **3** and MNM **4**.

Overlay of ψms23 with MNM-BODIPY showed that light-activated MNM **3** co-localized or tightly associated with *M. smegmatis* compared to non-activated MNM **3** (Figure 4 b, c, e and f). This is seen by the change in color of *M. smegmatis* from red to a lighter color in the overlay image, representing co-localization (Figure 5 f). In contrast, co-localization was absent or occurred at low levels in MNM **4** and non-activated MNM **3** (Figure 5 c, i, l). Fluorescent intensity was also significantly higher in light-activated MNM 3 compared to MNM **4** (p-value = 0.0123) (Figure 5 t). The increase in colocalization indicates that light-activated MNM **3** is more tightly associated with *M. smegmatis* either through embedding into the bacterial cell surface or entering into the bacteria.

To confirm that MNM with attached fluorophore BODIPY (MNM **3**) had nanomechanical properties similar to MNM **1**, we carried out viability assays with conditions used for confocal microscopy. Light-activated MNM **3** showed a 35% viability reduction when compared to a control without MNM, and a 30% viability reduction compared to MNM **4** (p-value = 0.0003 and 0.0016 respectively) (Figure 5 u, Supplemental Figure 3). In addition to colony forming unit (CFU) counts, we also quantified the amount of fluorescent emitted by *M. smegmatis* colonies exposed to MNM **3**, MNM **4,** and no MNM control (Supplemental Figure 4). We observe a significant reduction in fluorescent signal in *M. smegmatis* exposed to light-activated MNM **3** compared to MNM4 and no MNM control (p-value = 0.0013 and 0.0013 respectively) (Supplemental Figure 5). The reduction in florescent signals observed correlates with the reduction in bacterial viability observed (Figure 5 u). These results confirm that not only MNM **1**, but MNM **3** with large attachments (BODIPY fluorophore) on its stationary component still retains its nanomechanical antimicrobial properties. Our observations of confocal microscopy together with viability assays show that fast motor MNMs co-localize and tightly associate with *M. smegmatis* and cause a reduction in viability through its nanomechanical action on the bacteria. This imaging study provides a closer look at the mechanism of light-activated MNMs and adds to our understanding of how these MNMs carry out their nondrilling function on bacteria.

### Model illustrating the nanomechanical antimicrobial action of MNMs

Our results show that fast motor MNM **1** and **3** when activated with 365 nm light, cause a reduction in bacterial viability in *M. smegmatis, M. tuberculosis, S. aureus* and *E. coli*. This contrasts with the slow motor MNM **2** and **4** that did not show a significant reduction in bacterial viability with the same light activation. The confocal imaging study showed that MNM **3** co- localized with *M. smegmatis* with light-activation. Taking these observations together, we propose that upon light activation, the fast motor MNMs can embed and drill into bacterial cell walls through their nanomechanical action. This action causes disruptions in the bacteria cell walls resulting in the killing of the *M. smegmatis* bacterium. In contrast, slow motor MNMs are not able to penetrate the bacteria cell surface and do not have any bactericidal effects on the bacteria. (Figure 6).

**Figure 6.**
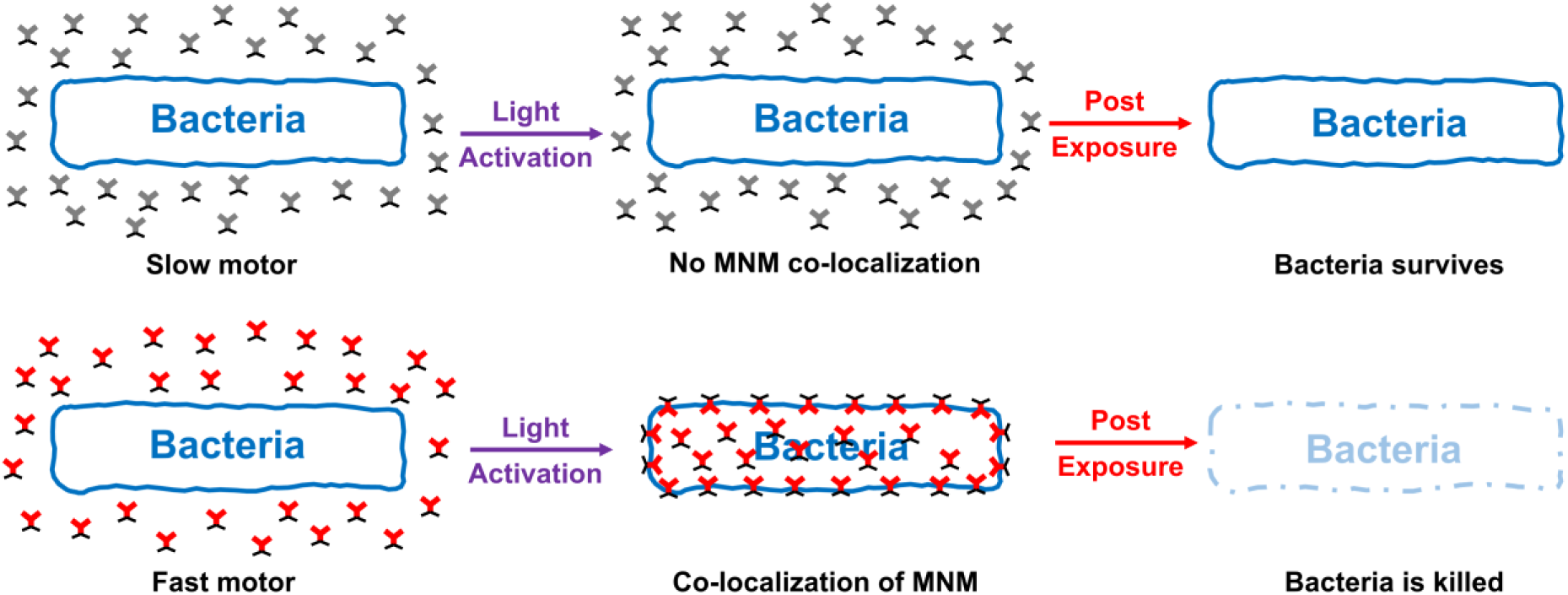
Model illustrating nanomechanical action of light-activated MNMs on bacteria. Bacteria exposed to MNMs have two distinct outcomes when activated by 365 nm light. Slow motors (MNM **2** and **4)** have no rotational action, remains outside the bacteria and have little to no effect on bacterial viability. Whereas fast motors (MNM **1**, **3,** and **5)** co-localize and embed into the bacterial cell wall causing disruptions that lead to a significant reduction in bacterial viability.

## Discussion

AR is an increasing global health crisis with far-reaching implications for public health, healthcare systems, and socio-economic factors (1, 27). The consequences of AR are multifaceted, encompassing increased mortality rates, compromised treatment efficacy, economic burdens, and challenges in the control of infectious diseases. Considering these consequences, concerted efforts are essential to address antimicrobial resistance comprehensively. AR is part of genomic diversity in bacterial species. The antibiotic resistome is found to be widespread and present in even ancient permafrost environmental specimens at a low occurrence rate (28, 29). However, with the wide use and misuse of present-day antibiotics, the frequency of AMR pathogens is increasing at a rapid pace and correlates with an increase in mortality among patients with otherwise treatable infections (3, 30). Conventional antibiotics favor the selection of AMR pathogens to the point that studies have shown that over 40% of the nosocomial *S. aureus* are now methicillin-resistant (MRSA) or multidrug-resistant (MDR) (31). Resistance to last-resort antibiotics such as carbapenems, among gram-negative bacteria in clinical settings occurs at an alarming rate requiring new solutions for this growing problem (31–33).

Recent studies by our collaborative research team shifts paradigms in antimicrobial treatment from previous biological approaches to a nanomechanical approach that may have a greater potential to be an effective antimicrobial agent based on the unique structure and novel mechanism of action of the nanomachine (13, 14). This study results show that light-activated fast MNMs acts as potent antimicrobial agents with as much as 97% reduction in bacterial viability in about 30 minutes of treatment. Unlike conventional antibiotics, it is unlikely for pathogens to develop resistance to a mechanical form of an antimicrobial agent, as the nanomachine did not need a specific target to act on. While it takes a long period of time to form AR, pathogenic bacteria were shown to not form any resistance to MNMs in over 20 bacterial growth cycles (14).

Our study adds to our current understanding of the function of light-activated nanomachines by showing that they co-localize with the bacteria upon activation (Figure 6). MNMs are likely to first localize close to the bacteria and when activated with light energy, embed into the bacterial cell wall by their nanomechanical action, MNM causes structural damage to the integrity of the bacterial surface. We showed this co-localization and internalization of MNM **3** with *M. smegmatis* upon light activation (Figures 5 and 6) by fluorescence confocal microscopy.

There are some limitations of this study. First, we use 365 nm light which is in the range of UV-c and it can have some harmful effects on living tissue. This limited the exposure time we could use to activate the MNM. Of the bacteria used in this study, *M. tuberculosis* was especially sensitive to 365 nm light activation over 5 minutes. We predict that with the increase of MNM activation time, the antimicrobial effect of MNM may increase several-fold as we saw with the 395 nm activated MNM **5**. The sensitivity to 365 nm light is expected because *M. tuberculosis* is not an environmental *Mycobacterium* and does not produce the pigments that protect environmental mycobacteria from UV (sun) light (34). Therefore, UV light is used in clinical or environmental settings to control to spread of M. tuberculosis and other pathogenic mycobacteria that do not provide protective pigments (23–25). This explains our observations that *M. tuberculosis* was much more sensitive to 365 nm light compared to *M. smegmatis*.

Second, 365 nm light has a relatively low penetration in host organs and tissue. This limits the effectiveness of MNM for the treatment of deep tissue infections. However, with our current approach, MNM can be potentially used as an antimicrobial agent to treat local infections including skin, mouth, lung, and urinary tract infections. In each of these target organs, MNM can be activated with light from an external source. However, to address both the above concerns, our group is currently working on the next generation of MNM that can be activated by visible light and near-infra-red (NIR) light. This will greatly increase the ability of MNM activation in much deeper host targets and also allow the activation of MNM for a longer time to achieve a far superior antimicrobial efficacy, without any harmful effects on the host.

Third, in this study, we used MNM which was only the motor component, without any specific binding affinity to the bacteria. The lack of specificity towards target bacteria may also explain the relatively low reduction in viability that we observed. When MNM 1 was used to target and permeabilize cancer cells, they had short-sequence peptides that allowed selective binding and high specificity (35). Targeting specific bacterial receptors can be achieved by selecting specific ligands on the complex cell wall structure in bacteria (36). Studies have shown that globotriaosylceramide (GB3) and lactosylceramide ligands bind LecA receptors with specificity in *Pseudomonas aeruginosa* and can be used for drug delivery in these pathogens (37, 38). We propose that these MNMs could carry pathogen-specific ligands on their stationary component and bind specifically to any pathogen of interest. Having more MNM closely attached to the bacteria allows its nanomechanical damage on the bacterial surface to be very effective; thereby increasing its efficacy as a nanomechanical antimicrobial agent.

In this study, we provide evidence that light-activated fast motor MNMs can reduce bacterial viability, particularly in mycobacteria setting them up as innovative agent to address the increase in antibiotic resistance. Our investigation into the nanomechanical action of light-activated MNMs on *M. smegmatis* provides valuable insights into combating antibiotic resistance in mycobacterial infections. The demonstrated antibacterial efficacy of fast motor MNMs, particularly MNM **1** and **3**, against *M. smegmatis*, *M. tuberculosis*, *S. aureus*, and *E. coli* underscores their potential as broad-spectrum antimicrobial agents. These findings hold clinical implications for addressing the growing challenges of antibiotic resistance in mycobacterial infections. In particular, the promise of MNMs for the treatment of MDR, XDR, and TDR *M. tuberculosis* strains is supported by our observation that the level of killing observed in *M. smegmatis* translates directly to *M. tuberculosis*. MNMs, with their unique mechanism of action and demonstrated effectiveness across diverse bacterial species, offer a promising avenue for future antimicrobial interventions. One might argue that treatment of tuberculosis could not easily be accomplished using MNMs due to the wavelength of light needed and the inability of that light to penetrate deep lung tissues where *M. tuberculosis* is known to reside. However, the recent development of long-wavelength MNMs demonstrates that such roadblocks could be overcome with the continued development of MNMs that combine mycobactericidal activity and long-wavelength activation (39). The nanomechanical action of the MNM makes it a unique, effective, and potentially potent antimicrobial agent that can help us in the battle against multidrug-resistant pathogens.

## Supplemental Information

Extended experimental procedures and supplementary figures are included in the Supplemental Information and can be found separately.

## Acknowledgments

The work at Rice University was funded by the Discovery Institute and the Welch Foundation (G10000837). RSG was supported by the Discovery Institute and the Roth Family. JDC was supported in part by NIH grant AI186092 and AI149383

## References

1. Global burden of bacterial antimicrobial resistance in 2019: a systematic analysis. Lancet. 2022;399(10325):629-55.

2. Salam MA, Al-Amin MY, Salam MT, Pawar JS, Akhter N, Rabaan AA, et al. Antimicrobial Resistance: A Growing Serious Threat for Global Public Health. Healthcare (Basel). 2023;11(13).

3. O’Neill J. Tackling Drug-Resistant Infections Globally: Final Report and Recommendations. Review on Antimicrobial Resistance. 2016:1–84.

4. Dheda K, Gumbo T, Gandhi NR, Murray M, Theron G, Udwadia Z, et al. Global control of tuberculosis: from extensively drug-resistant to untreatable tuberculosis. Lancet Respir Med. 2014;2(4):321–38.

5. Loiseau C, Windels EM, Gygli SM, Jugheli L, Maghradze N, Brites D, et al. The relative transmission fitness of multidrug-resistant Mycobacterium tuberculosis in a drug resistance hotspot. Nat Commun. 2023;14(1):1988.

6. Saxena S, Spaink HP, Forn-Cuní G. Drug Resistance in Nontuberculous Mycobacteria: Mechanisms and Models. Biology (Basel). 2021;10(2).

7. Seung KJ, Keshavjee S, Rich ML. Multidrug-Resistant Tuberculosis and Extensively Drug-Resistant Tuberculosis. Cold Spring Harb Perspect Med. 2015;5(9):a017863.

8. Parida SK, Axelsson-Robertson R, Rao MV, Singh N, Master I, Lutckii A, et al. Totally drug-resistant tuberculosis and adjunct therapies. J Intern Med. 2015;277(4):388–405.

9. Sachan RSK, Mistry V, Dholaria M, Rana A, Devgon I, Ali I, et al. Overcoming Mycobacterium tuberculosis Drug Resistance: Novel Medications and Repositioning Strategies. ACS Omega. 2023;8(36):32244–57.

10. Udwadia ZF. MDR, XDR, TDR tuberculosis: ominous progression. Thorax. 2012;67(4):286–8.

11. Brown ED, Wright GD. Antibacterial Drug Discovery in the Resistance Era. Nature. 2016;529:336–43.

12. Palomino JC, Martin A. Drug Resistance Mechanisms in Mycobacterium tuberculosis. Antibiotics (Basel). 2014;3(3):317–40.

13. Galbadage T, Liu D, Alemany LB, Pal R, Tour JM, Gunasekera RS, et al. Molecular Nanomachines Disrupt Bacterial Cell Wall, Increasing Sensitivity of Extensively Drug-Resistant Klebsiella pneumoniae to Meropenem. ACS nano. 2019;13:14377–87.

14. Santos AL, Liu D, Reed AK, Wyderka AM, van Venrooy A, Li JT, et al. Light-activated molecular machines are fast-acting broad-spectrum antibacterials that target the membrane. Sci Adv. 2022;8(22):eabm2055.

15. García-López V, Chiang P-T, Chen F, Ruan G, Martí AA, Kolomeisky AB, et al. Unimolecular Submersible Nanomachines. Synthesis, Actuation, and Monitoring. Nano Lett. 2015;15:8229–39.

16. Garcia-Lopez V, Chen F, Nilewski LG, Duret G, Aliyan A, Kolomeisky AB, et al. Molecular Machines Open Cell Membranes. Nature. 2017;548:567–72.

17. Liu D, Garcia-Lopez V, Gunasekera RS, Greer Nilewski L, Alemany LB, Aliyan A, et al. Near-Infrared Light Activates Molecular Nanomachines to Drill into and Kill Cells. ACS Nano. 2019;13:6813–23.

18. Gunasekera RS, Galbadage T, Ayala-Orozco C, Liu D, García-López V, Troutman BE, et al. Molecular Nanomachines Can Destroy Tissue or Kill Multicellular Eukaryotes. ACS Appl Mater Interfaces. 2020;12(12):13657–70.

19. Bansal-Mutalik R, Nikaido H. Mycobacterial outer membrane is a lipid bilayer and the inner membrane is unusually rich in diacyl phosphatidylinositol dimannosides. Proceedings of the National Academy of Sciences. 2014;111(13):4958–63.

20. Sparks IL, Derbyshire KM, Jacobs WR, Jr., Morita YS. Mycobacterium smegmatis: The Vanguard of Mycobacterial Research. J Bacteriol. 2023;205(1):e0033722.

21. Chakaya J, Khan M, Ntoumi F, Aklillu E, Fatima R, Mwaba P, et al. Global Tuberculosis Report 2020 -Reflections on the Global TB burden, treatment and prevention efforts. Int J Infect Dis. 2021;113 Suppl 1(Suppl 1):S7-s12.

22. Gygli SM, Borrell S, Trauner A, Gagneux S. Antimicrobial resistance in Mycobacterium tuberculosis: mechanistic and evolutionary perspectives. FEMS Microbiol Rev. 2017;41(3):354–73.

23. Xu P, Peccia J, Fabian P, Martyny JW, Fennelly KP, Hernandez M, et al. Efficacy of ultraviolet germicidal irradiation of upper-room air in inactivating airborne bacterial spores and mycobacteria in full-scale studies. Atmospheric Environment. 2003;37(3):405–19.

24. Escombe AR, Moore DA, Gilman RH, Navincopa M, Ticona E, Mitchell B, et al. Upper-room ultraviolet light and negative air ionization to prevent tuberculosis transmission. PLoS Med. 2009;6(3):e43.

25. Shleeva M, Savitsky A, Kaprelyants A. Photoinactivation of mycobacteria to combat infection diseases: current state and perspectives. Appl Microbiol Biotechnol. 2021;105(10):4099–109.

26. Batt SM, Minnikin DE, Besra GS. The thick waxy coat of mycobacteria, a protective layer against antibiotics and the host’s immune system. Biochem J. 2020;477(10):1983–2006.

27. Dadgostar P. Antimicrobial Resistance: Implications and Costs. Infect Drug Resist. 2019;12:3903–10.

28. Perry JA, Westman EL, Wright GD. The antibiotic resistome: what’s new? Curr Opin Microbiol. 2014;21:45–50.

29. Forsberg KJ, Patel S, Gibson MK, Lauber CL, Knight R, Fierer N, et al. Bacterial phylogeny structures soil resistomes across habitats. Nature. 2014;509(7502):612-6.

30. Cosgrove SE, Sakoulas G, Perencevich EN, Schwaber MJ, Karchmer AW, Carmeli Y. Comparison of mortality associated with methicillin-resistant and methicillin-susceptible Staphylococcus aureus bacteremia: a meta-analysis. Clin Infect Dis. 2003;36(1):53–9.

31. Weinstein RA. Controlling antimicrobial resistance in hospitals: infection control and use of antibiotics. Emerg Infect Dis. 2001;7(2):188–92.

32. Nordmann P, Poirel L. Emerging carbapenemases in Gram-negative aerobes. Clin Microbiol Infect. 2002;8(6):321–31.

33. Livermore DM, Woodford N. Carbapenemases: a problem in waiting? Curr Opin Microbiol. 2000;3(5):489–95.

34. Tran T, Dawrs SN, Norton GJ, Virdi R, Honda JR. Brought to you courtesy of the red, white, and blue-pigments of nontuberculous mycobacteria. AIMS Microbiol. 2020;6(4):434–50.

35. Garcia-Lopez V, Chen F, Nilewski LG, Duret G, Aliyan A, Kolomeisky AB, et al. Molecular machines open cell membranes. Nature. 2017;548(7669):567-72.

36. Brown L, Wolf JM, Prados-Rosales R, Casadevall A. Through the wall: extracellular vesicles in Gram-positive bacteria, mycobacteria and fungi. Nature reviews Microbiology. 2015;13(10):620–30.

37. Muller SK, Wilhelm I, Schubert T, Zittlau K, Imberty A, Madl J, et al. Gb3-binding lectins as potential carriers for transcellular drug delivery. Expert Opin Drug Deliv. 2017;14(2):141–53.

38. Zheng S, Eierhoff T, Aigal S, Brandel A, Thuenauer R, de Bentzmann S, et al. The Pseudomonas aeruginosa lectin LecA triggers host cell signalling by glycosphingolipid-dependent phosphorylation of the adaptor protein CrkII. Biochim Biophys Acta. 2017;1864(7):1236–45.

39. Ayala-Orozco C, Galvez-Aranda D, Corona A, Seminario JM, Rangel R, Myers JN, et al. Molecular jackhammers eradicate cancer cells by vibronic-driven action. Nat Chem. 2023.

