## Supplemental Material for "Microbicidal Mechanisms for Light-Activated Molecular Nanomachines in *Mycobacterium smegmatis*: A Model for Pathogenic Bacteria"

*Thushara Galbadage[1], Dongdong Liu[3], James M. Tour[3,4,5,6], Jeffrey D. Cirillo[1,2*], and Richard S. Gunasekera[3,7,8*]*

*[1]Department of Microbial Pathogenesis and Immunology, Texas A&M University, School of Medicine, [2]Center for Airborne Pathogens Research and Imaging, Texas A&M Institute for Genome Sciences and Society, Bryan, TX*

*[3]Department of Chemistry, [4]Department of Materials Science and NanoEngineering, [5]Smalley-Curl Institute, [6]NanoCarbon Center, [7]Department of BioSciences, Rice University, Houston, TX*

*[8]Department of Biological Science, Biola University, La Mirada, CA*

*Corresponding authors; authors contributed equally

**Keywords**: nanomolecules, molecular machines, multidrug-resistant, mycobacteria, *M. tuberculosis*, *M. smegmatis*, *S. aureus*, *E. coli*

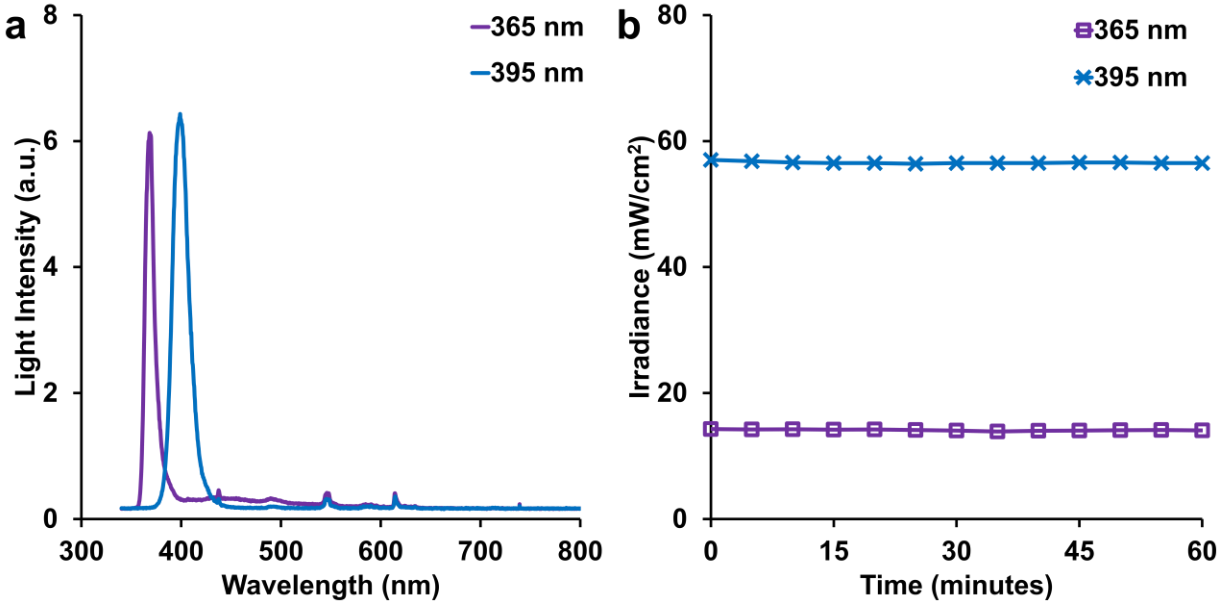


**Supplemental Figure 1. The emission spectra of the 365 nm and 395 nm light sources.** A spectrum test was performed on the 365 nm and 395 light sources used in this study, using a spectrograph. The 365 nm light source had an emission spectrum from about 358 to 386 nm, with a peak intensity at 368 nm wavelength. The 395 nm light source had an emission spectrum from about 382 to 422 nm, with a peak intensity at 398 nm wavelength.

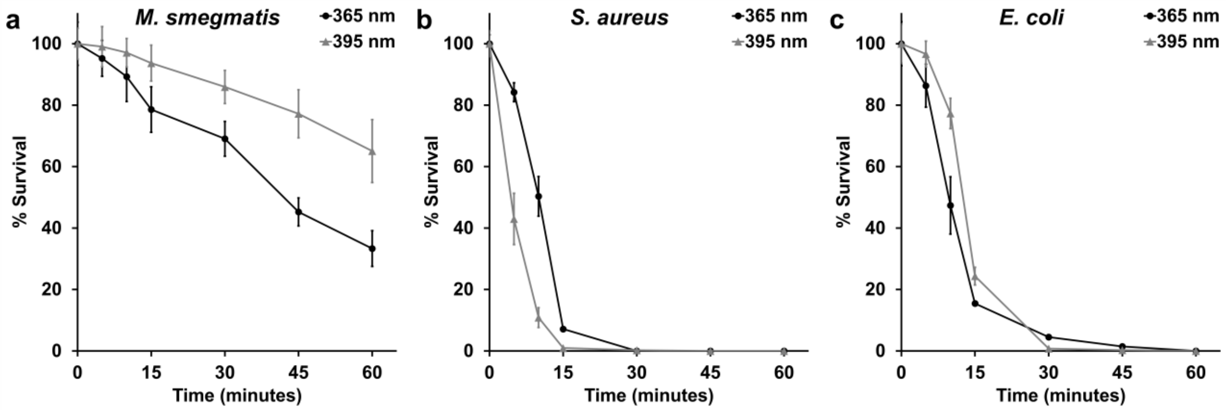


**Supplemental Figure 2.** Bactericidal effect of the 365 nm and 395 nm light sources on *M. smegmatis*, *S. aureus,* and *E. coli* over 60 minutes of constant light exposure.

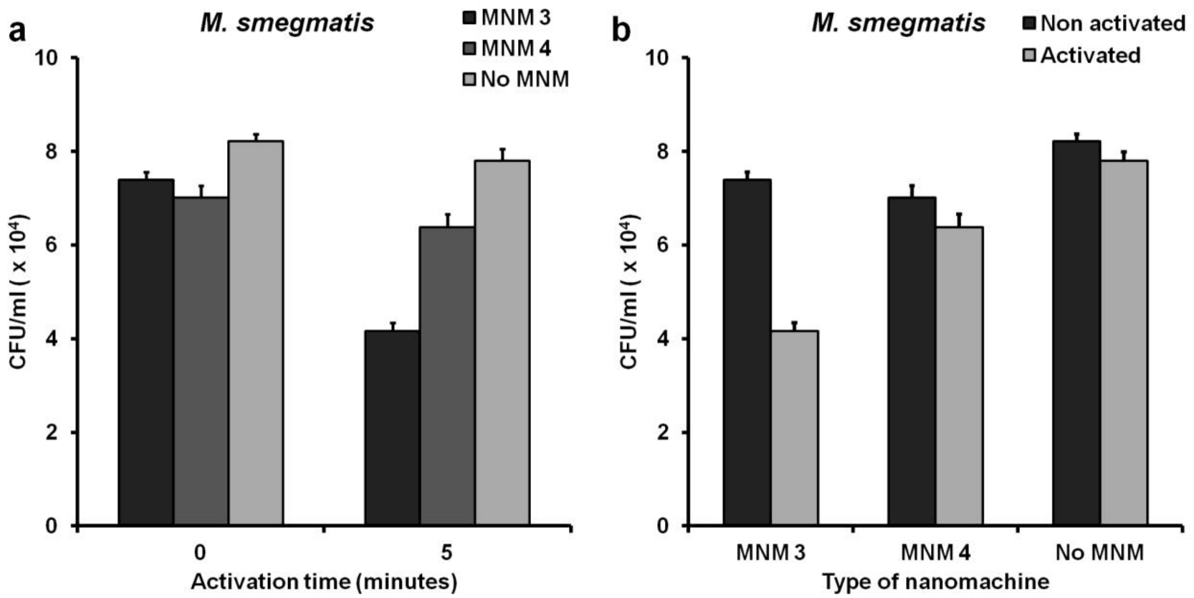


**Supplemental Figure 3 Viability in CFU/ml of *M. smegmatis* (ψms23) in Activated to Non-Activated MNM**

tdTomato expressing *M. smegmatis* (ψms23) exposed to 10 μM of MNM. (a) Comparison of ψms23 CFU/ml in MNM **1**, MNM **2,** and No MNM groups. (b) Comparison of ψms23 CFU/ml with and without light activation.

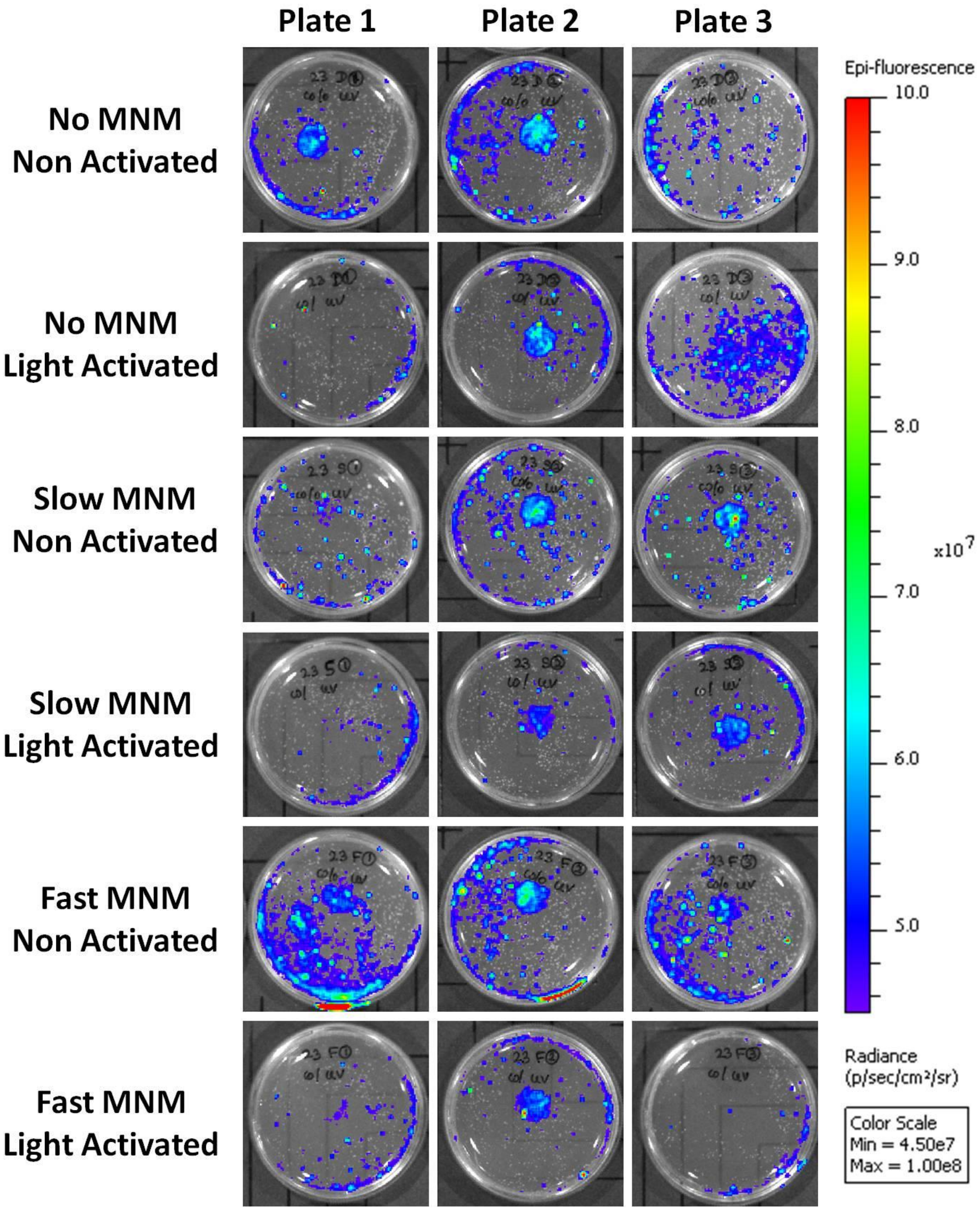


**Supplemental Figure 4 IVIS Image of tdTomato Fluorescent**

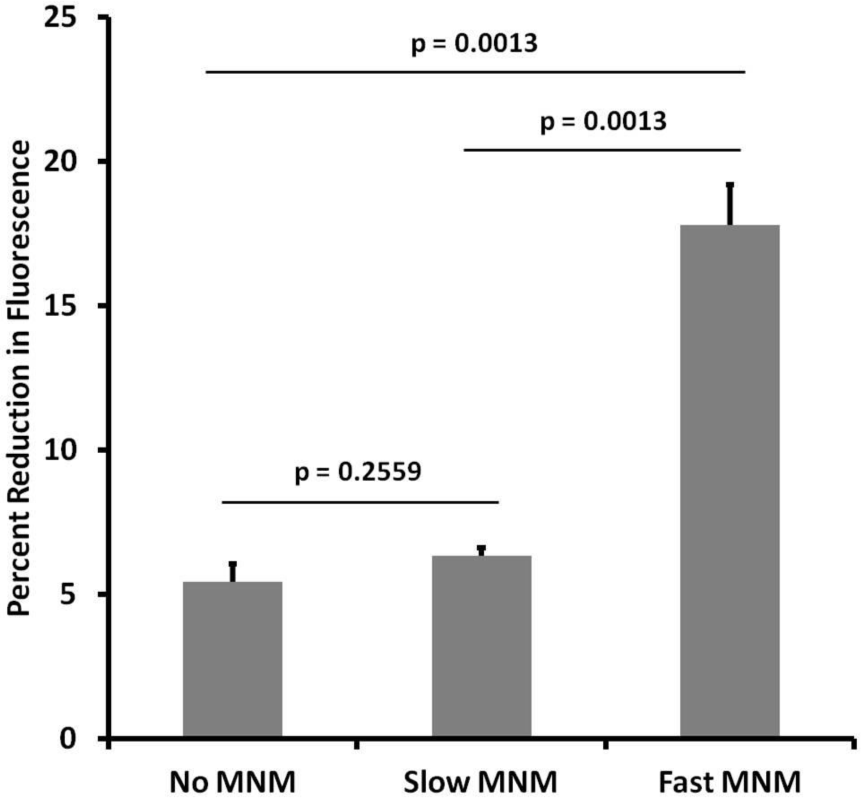


**Supplemental Figure 5 Relative Reduction in *M. smegmatis* (ψms23) in Activated to Non-Activated MNM**
